# Auditory timing precision is determined by spectro-temporal continuity across sounds, not individual sound acoustics

**DOI:** 10.64898/2026.09.26.754418

**Authors:** Charlotte M. Mock, Leon Ilge, Yulia Oganian

**Affiliations:** Centre for Integrative Neuroscience Tübingen; Graduate Training Centre of Neuroscience Tübingen; International Max Planck Research School for The Mechanisms of Mental Function and Dysfunction; University of Tübingen

## Abstract

Predicting when an event will occur is key to perceiving and evaluating dynamic sensory inputs, such as speech. The utility of temporal predictions hinges on perceptual timing precision (PTP), the sensitivity to deviations between actual and predicted sound onset times. Past work suggests that PTP is primarily determined by acoustic properties of target sounds, regardless of the acoustics of preceding sound cues. In contrast, in four experiments, we discover that PTP is best when context and target sound spectra and temporal envelopes match, with only minor effects of individual sound acoustics. This ability to utilize contextual continuity improved with musical aptitude. Theoretical modeling showed that spectrally tuned oscillatory entrainment to sound onsets and offsets is necessary to account for our results. In sum, the precision of temporal predictions is defined not by the acoustics of individual sounds but by the continuity of spectral and temporal content within a sound stream.

## Introduction

Sensory environments are inherently structured in time. Within this temporal structure, we are able to perceive when events occur, to track events over time, and to judge their relative timing ^1–3^. The timing of individual auditory events carries important information, especially for dynamic sensory inputs, such as speech and music ^4–7^. For example, a pianist hitting a key off beat or a deliberate speech pause are immediately noticeable, reflecting sensitivity to the (mis)alignment between actual and expected event timing. Such judgments hinge on perceptual timing precision (PTP), the sensitivity of the perceptual system to small deviations in event timing. High PTP may support perception by enabling precise predictions about the timing of upcoming sensory events. Conversely, even when temporal information is available, reduced PTP may limit the precision of such predictions, thereby constraining the utility of temporal information for perception.

Prior work suggests that PTP is determined by the temporal precision afforded by a sound ^8^, but also that PTP is enhanced with higher musical aptitude ^9–13^. Listeners are highly sensitive to temporal deviations in rhythmic sequences of impulse-like click sounds ^8,14^. However, speech and music sounds have a psycho-acoustically more complex spectro-temporal structure. In this case, a range of acoustic determinants defines the perceived temporal location ^15–20^. Crucially, even professional musicians perceive a range of physical onset times of such temporally extended sounds as aligned with the beat of an isochronous cue sequence^9^. It was found that the width of this range, termed a sound’s “beat bin” ^21^, depended on the target sound’s acoustic characteristics: for short sounds with fast-rising onsets (e.g., drum sounds) beat bins were narrower, indicating greater PTP, than for longer sounds with slow-rising onsets^8,14^.

A key potential implication of these findings is that speech, where individual units (e.g., syllables) are extended in time and have gradual onsets, should afford rather low PTP. However, inter-individual agreement in perceived event onsets within continuous speech is high ^22^, consistently anchored to salient acoustic-linguistic landmarks, such as vowel onsets. Moreover, motor synchronization to speech exhibits hallmark patterns closely resembling those observed with click sounds and music, such as stable phase locking and anticipatory timing of movements to the perceived beat ^23,24^. This suggests that temporal precision for speech and non-speech sounds relies on principally the same computational mechanisms. Together, these findings show that high PTP is possible for psycho-acoustically complex sounds, such as speech, and suggest that perceptual timing precision cannot be determined by target sound properties alone.

One hypothesis that can reconcile these findings is that PTP might additionally be influenced by the acoustic context. For example, in rhythmic streams, temporal prediction automatically engages oscillatory entrainment (phase alignment) to the external rhythm ^25–30^, which can then support beat deviation judgments. Recordings from macaque primary auditory cortex show that attention to rhythmic auditory stimuli entrains oscillations across tonotopically organized regions in a frequency-specific manner ^31^. This work found that neuronal populations tuned to the attended frequency align their high-excitability phases to expected stimulus timing, while populations tuned to non-attended frequencies align opposite, low-excitability phases. More recently, L’Hermité and Zoefel ^32^ found that rhythmic fluctuations in auditory target-detection sensitivity persisted after the offset of a rhythmic tone sequence. Importantly, the timing of optimal target detection depended on the frequency match between entraining and target sounds, providing a behavioral counterpart to the neural recordings reported by Lakatos and colleagues. Together, this raises the possibility that observations of reduced PTP for psycho-acoustically complex sounds may have been due to spectral mismatch with the preceding context. However, whether and how this extends to explicit temporal judgements and perceptual sensitivity to temporal deviations remains unknown.

To close this gap, across four experiments, we tested whether beat bin width, with narrower bins indicating greater PTP, reflects an intrinsic psycho-acoustic property of individual sounds or instead depends on contextual factors. We implemented an iterative adjustment paradigm, where participants repeatedly estimated the temporal offset between a target sound and a preceding rhythmic cue sequence, yielding individual PTP estimates for different cue-target sound combinations. This paradigm allowed us to independently manipulate the acoustic properties of cue and target sounds and assess their unique contributions to PTP across individuals varying in musical aptitude. Surprisingly, we found that, in an acoustically matching context, PTP was as good for psycho-acoustically complex sounds as for simple click sounds. The key determinant of PTP was the spectral match between cue and target sounds, with additional contributions of matching temporal envelopes in musically apt participants. In contrast, refuting prior theories, target-sound acoustics did not affect PTP. Computational modeling of PTP in rhythmic contexts based on frequency-specific oscillatory entrainment captured all key observed behavioral patterns, including musicality effects. Overall, we reveal hitherto unknown determinants of PTP in dynamic contexts and offer a mechanistic framework for future evaluation of perceptual timing across a broad range of rhythmic sensory contexts.

## Results

To assess the effects of different cue and target sounds on individuals’ perceptual timing precision (PTP), we implemented the iterative adjustment paradigm (IAP; adapted from ^16^). Participants listened to isochronous cue sound sequences, followed by a target sound, initially presented off-beat relative to the expected next beat of the cue sequence (**Fig. 1A**). Participants iteratively adjusted the target sound’s timing relative to the cue sequence on a visual analog scale, until they perceived the target as on beat (**Fig. 1B**). We then estimated PTP for each participant and cue-target combination as the standard deviation of target asynchronies after final adjustments.

**Fig. 1.**
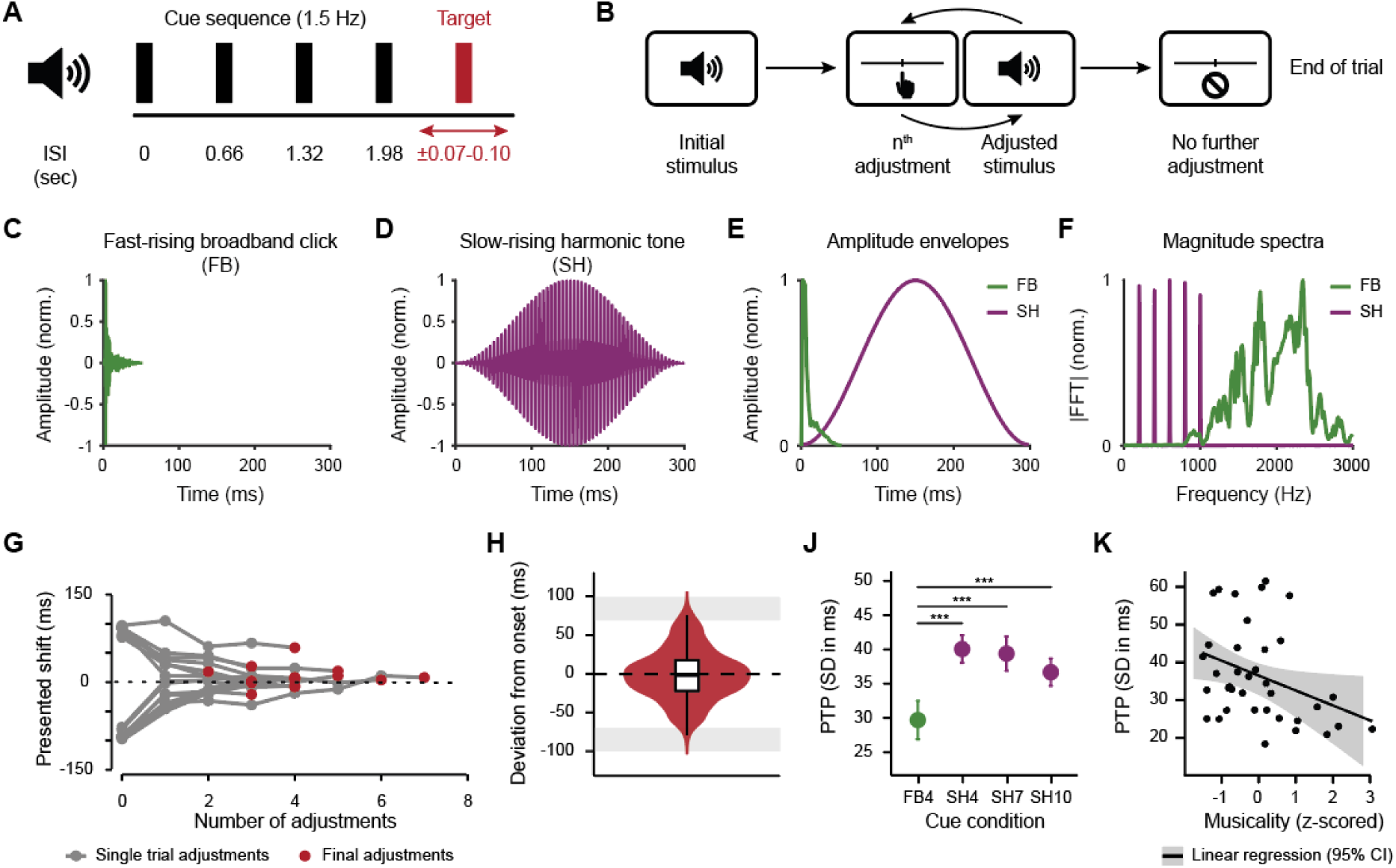
Iterative adjustment paradigm and Experiment 1. **A.** Cue-target sequences in Experiment 1 consisted of four to seven cues (black) presented in isochrony at a rate of 1.5 Hz and a single target sound (red) occurring at an offset of ±70-100 ms. In Exp. 2-3, the cue sequence always contained four cues. **B.** Schematic description of the iterative adjustment paradigm (IAP). First, participants listen to a cue-target sequence; next, they are presented with a horizontal slider on the screen with which they can adjust the temporal position of the target sound. Then, they are presented with the adjusted version of the sequence. This listen-adjust-listen cycle is repeated until participants perceive the entire sequence as isochronous, or until a maximum of 12 adjustments is reached. **C-D.** Waveforms of stimulus sounds used in Experiments 1 and 2a+b. **E.** Overlaid amplitude envelopes of stimuli. **F.** Overlaid normalized magnitude spectra of stimuli. **G.** Exemplary adjustment trajectories of one participant in the baseline (FB4) condition in Experiment 1. **H.** Group-level distribution of temporal deviations between final adjusted target-sound onsets and the acoustic onset of the expected next beat in the baseline condition (FB4) reflects target positions perceived as isochronous with the preceding cues. Positive values indicate target onsets occurring after the expected beat, negative values indicate target onsets occurring before the expected beat. Shaded bands mark the initial target-offset ranges of ±70–100 ms. **J.** Perceptual timing precision (PTP) in Experiment 1. Mean ±1 SEM, *** p < .005. PTP was best, that is, SD was lowest, for sequences with FB cues (green), whereas increasing the number of SH cues (purple) did not improve PTP. **K.** Across task conditions in Experiment 1, higher musicality was associated with better PTP.

### Experiment 1: Perceptual timing precision does not improve with cue sequence length

We tested whether PTP improves with context length in psycho-acoustically complex contexts, e.g., when the context consists of long harmonic sounds with a slow-rising onset, characteristic of speech and music. Longer cue sequences may potentially allow listeners to form more precise temporal expectations despite acoustic complexity.

In all conditions, the target was a short click-like sound (fast-rising broadband click, FB, 52 ms duration; **Fig. 1C-F**). In the baseline condition (FB4), it was preceded by four identical FB sounds. In the complex cue conditions, 4, 7, or 10 complex harmonic sound cues preceded the same click target (SH4, SH7, SH10). The SH sound was a slow-rising harmonic tone, consisting of five harmonics starting at a fundamental frequency of 200 Hz, with a symmetric rise and decay of 150 ms and an overall duration of 300 ms (**Fig. 1D-F**). The initial position of the target on each trial was jittered relative to the expected next beat of the isochronous cue sequence, with initial asynchronies ranging from ±70 to ±100ms.

To verify that participants (n=36) performed the IAP as intended, we first examined adjustment behavior in the FB4 condition. Participants made on average 2.76 (SD = 1.77) adjustments per trial, reducing the initial offsets to a final offset of −0.70 ms on average (SD = 7.77 ms; see **Fig. 1G** for exemplary adjustment trajectories). Consistent with successful adjustment behavior, final target positions in the baseline condition clustered around the next on-beat time (**Fig. 1H**).

We then asked how PTP changes with slow-rising harmonic cues. Linear mixed-effects modeling showed that PTP was worse for complex than for simple cues, irrespective of cue sequence length (comparison to FB4 condition for SH4, *β* = 10.19, 95% CI [6.17, 14.21], *SE* = 2.03, *t*(97.36) = 5.03, *p* < .001, ηp^2^ = .21; SH7, *β* = 9.71, 95% CI [5.77, 13.65], *SE* = 1.99, *t*(96.80) = 4.89, *p* < .001, ηp^2^ = .20; SH10, *β* = 8.07, 95% CI [3.88, 12.26], *SE* = 2.11, *t*(97.94) = 3.83, *p* < .001, ηp^2^ = .13; **Fig. 1J**), replicating previous reports ^14^. Critically, when cues were complex, PTP did not improve with cue sequence length (SH4 vs. SH7: *p* > .5, ηp^2^ < .01; SH4 vs. SH10: *p* = .2, ηp^2^ = .02). A Bayesian model comparison provided additional moderate evidence for the absence of a cue-length effect (BF_01_ = 5.73). Furthermore, while participants with higher musicality scores exhibited greater PTP overall (*β* = −5.01, 95% CI [−9.19, −0.84], *SE* = 2.09, *t*(62.60) = −2.40, *p* = .019, ηp^2^ = .08; **Fig. 1K**), this beneficial effect did not differ between conditions (all *p* ≥ .28).

Experiment 1 showed less precise perceptual timing for click targets in the context of long, slow-rising harmonic tones than in the context of click sounds, in line with prior research ^8,14^. Moreover, PTP did not improve when the cue sequence was extended ^33^. However, a potential confound of our and past experiments is that conditions differed not only in the acoustic structure of the cue sound, but also in spectro-temporal similarity between cue and target sounds. We thus hypothesized that PTP may have been lower for slow-rising than for fast-rising cues in Experiment 1, due to spectro-temporal differences between cue and target, rather than due to cue acoustics (**Fig. 2A** left panel). This stands in contrast to the dominant view in the literature that emphasizes the characteristics of individual sounds ^8,14^, predicting greater PTP when either cue or target sound have fast-rising onsets (**Fig. 2A** middle and right panels).

**Fig. 2.**
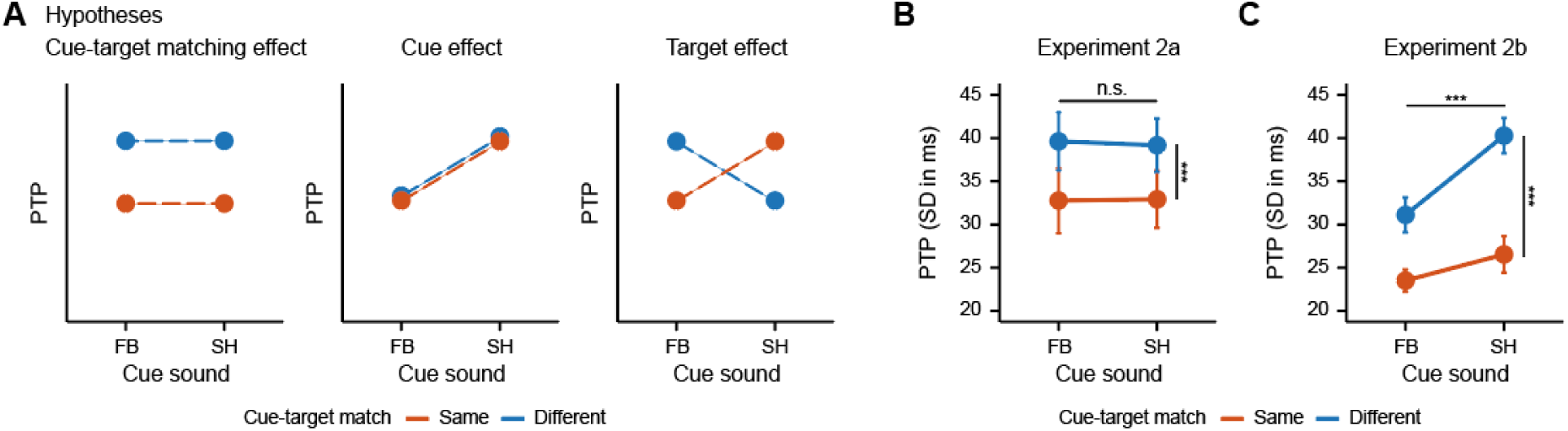
Experiment 2. **A.** Alternative hypotheses for Experiment 2. We hypothesized that a cue–target match effect (left), rather than effects of cue (middle) or target (right) acoustics might determine PTP. **B+C.** Perceptual timing precision (PTP) in Experiments 2a and 2b. Observed condition means ±1 SEM. n.s. p > 0.05, *** p < .005. FB, fast-rising broadband click; SH, slow-rising harmonic tone. **B.** In Experiment 2a, PTP was greatest when cue and target sounds were identical. There were no effects of cue or target sounds. **C.** In Experiment 2b, PTP was again greatest when cue and target were identical. Additionally, PTP was improved when targets were preceded by a FB cue than by a SH cue.

### Experiment 2: Cue-target continuity, not target sound properties determine PTP

To test this hypothesis, in Experiment 2a, participants (online, n=21) performed the IAP on sequences where both cue and target could be either the fast-rising broadband click (FB, **Fig. 1C**) or the slow-rising harmonic tone (SH, **Fig. 1D**) from Experiment 1. This resulted in an orthogonal manipulation of cue sound identity and the spectro-temporal match between cue and target across four conditions, presented in a blocked experimental design. As hypothesized, we found better PTP when cue and target were identical than when they differed (*β* = −4.28, 95% CI [−5.99, −2.57], *SE* = 0.85, *t*(54.03) = −5.01, *p* < .001, ηp^2^ = .32; **Fig. 2B**). In contrast, neither cue (*β* = 0.33, 95% CI [−1.38, 2.04], *SE* = 0.85, *t*(54.05) = 0.39, *p* = .701, ηp^2^ < .01; **Fig. 2B**) nor target acoustics (*β* = −0.36, 95% CI [−2.07, 1.36], *SE* = 0.85, *t*(54.05) = −0.42, *p* = .678, ηp^2^ < .01; **Fig. 2B**), nor musicality modulated PTP (all *p* ≥ .125).

Next, we replicated the experiment in a laboratory setting (Exp. 2b, n=21). Again, participants had more precise perceptual timing when cue and target were identical than when they differed (*β* = −5.01, 95% CI [−6.75, −3.27], *SE* = 0.87, *t*(49.75) = −5.79, *p* < .001, ηp^2^ = .40; **Fig. 2C**). Here, however, PTP was higher for FB cues than for SH cues (*β* = −3.19, 95% CI [−4.92, −1.45], *SE* = 0.86, *t*(49.25) = −3.69, *p* < .001, ηp^2^ = .22), whereas the effect of target sound was not significant (*p* > .1, ηp^2^ = .04). As in Experiment 2a, musicality did not affect PTP (all *p* ≥ .117).

In sum, Experiments 2a and 2b reveal that acoustic continuity between cue and target sounds determines PTP. Notably, we found no effect of target sound properties, calling into question the concept of beat bins as a context-independent property of sounds.

### Experiment 3. Distinct roles of spectral and temporal cue-target continuity in PTP

Next, to understand how spectral and temporal sound properties contribute to the continuity effect, we independently manipulated spectral content and temporal dynamics of stimulus sounds in Experiment 3. We hypothesized that cue-target continuity along each dimension might contribute to PTP. In addition, we tested whether these dimensions contribute independently or interactively. If their effects depend on alignment across dimensions, the greatest benefits should occur when cue and target are identical.

The cue in all four conditions was the SH sound used in Experiments 1 and 2 (**Fig. 3A-C**). Targets either had the same symmetric slow-rising envelope as the cue, or were short and fast-rising, with a slower decay trajectory (**Fig. 3A-B**). Further, targets either had the same harmonic spectrum as the cue or had a non-harmonic broadband spectrum (**Fig. 3C**), resulting in a 2 x 2 orthogonal manipulation of spectral and temporal envelope match. Participants (n=32) evaluated the four cue-target sequences in a blocked design in the IAP.

**Fig. 3.**
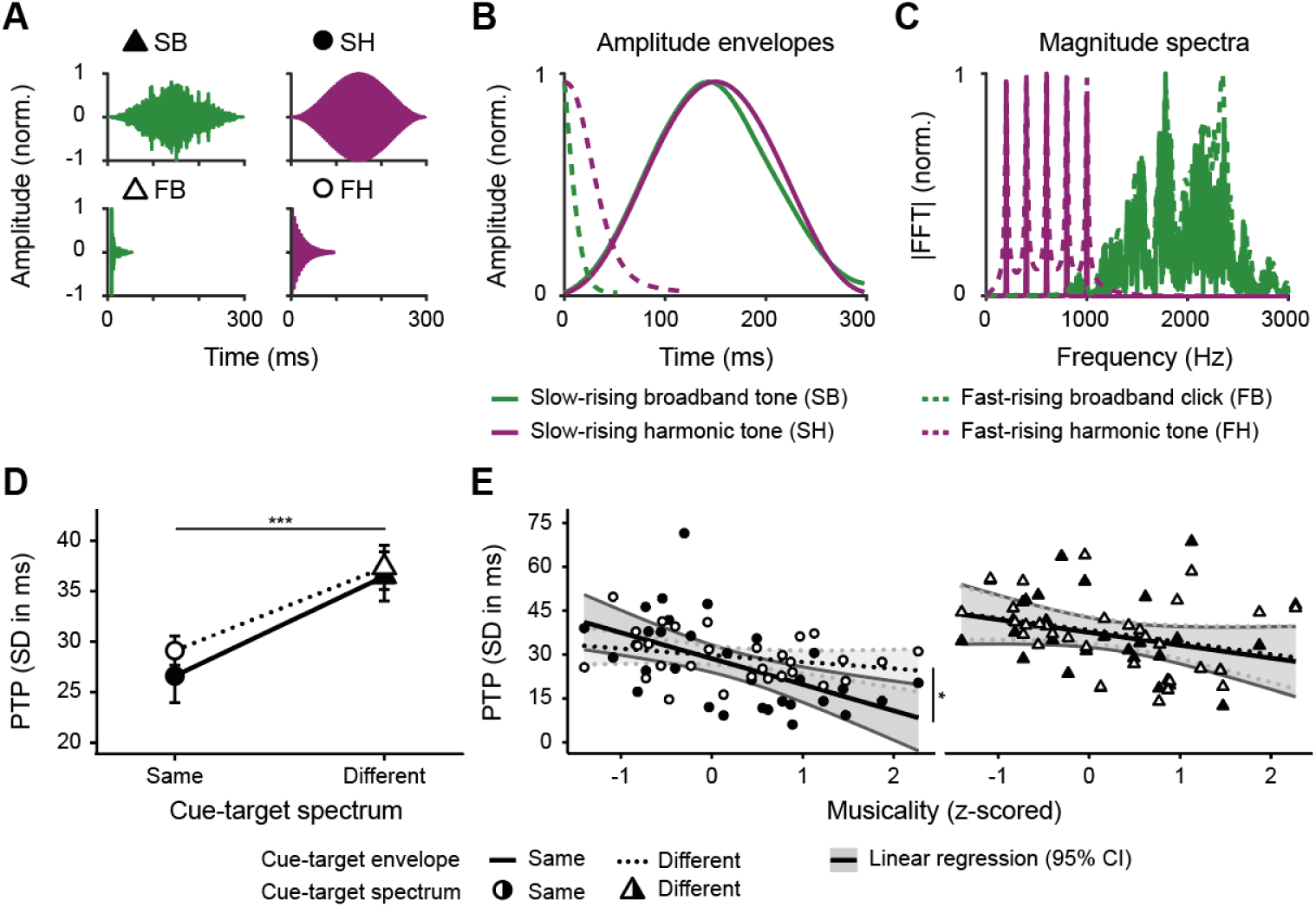
Experiment 3. **A.** Stimulus waveforms. SH and FB were identical to the sounds used in Experiments 1 and 2a+b. Sound SH was the cue in all conditions. **B.** Overlaid stimulus amplitude envelopes. **C.** Overlaid normalized stimulus magnitude spectra. **D.** PTP as a function of cue–target spectrum and envelope match. Observed condition means ±1 SEM. *** p < .005. PTP was best when cue and target sounds had the same spectrum. **E.** Higher musical aptitude was associated with greater PTP, with the strongest benefit from musicality when cue and target had identical spectrum and envelope. *p<0.05.

At the group level (**Fig. 3D)**, PTP was best when cue and target had the same spectrum (*β* = 9.21, 95% CI [5.34, 13.08], *SE* = 1.95, *t*(83.56) = 4.73, *p* < .001, ηp^2^ = .21) but was not influenced by envelope mismatch (*β* = 2.62, 95% CI [−1.36, 6.59], *SE* = 2.00, *t*(84.09) = 1.31, *p* > .194, ηp^2^ = .02). However, this pattern was modulated by musical aptitude (**Fig. 3E**). First, PTP for identical cue-target sequences improved with musicality (*β* = −8.88, 95% CI [−13.65, −4.10], *SE* = 2.38, *t*(52.27) = −3.73, *p* < .001, ηp^2^ = .21). Moreover, the beneficial effect of matching spectra between cues and target increased with musicality (*β* = 4.29, 95% CI [0.16, 8.42], *SE* = 2.08, *t*(83.47) = 2.07, *p* = .042, ηp^2^ = .05). Along the same lines, a benefit of matching envelopes emerged with musical aptitude (*β* = 5.66, 95% CI [1.49, 9.83], *SE* = 2.10, *t*(83.63) = 2.70, *p* = .008, ηp^2^ = .08). This effect was marginally stronger when cue and target had matching spectra (3-way interaction: *β* = –5.74, 95% CI [−11.61, 0.13], *SE* = 2.95, *t*(83.49) = –1.94, *p* = .055, ηp^2^ = .04). That is, musically apt participants benefitted additionally when both spectra and temporal envelopes of cue and target sounds were identical.

In sum, PTP for complex sounds is primarily determined by spectral continuity between cue and target sounds. Additionally, benefits from matching temporal envelopes emerged in individuals with higher musicality. That is, musicality not only generally enhances receptive timing abilities, but also specifically enhances listeners’ ability to leverage spectral patterns and temporal envelope dynamics for timing precision. These findings specify the importance of the acoustic context when evaluating temporal perception and prediction.

### Spectrally tuned oscillator model predicts cue-target continuity effects

We next sought to extend existing models of oscillatory entrainment to account for our results. The dominant approach to model neural entrainment to acoustic streams uses a single fixed-frequency oscillator, with temporal frequency corresponding to the average frequency of events in the cue stream ^26,34,35^. This model implements neural entrainment to the cue stream via gradual phase reset towards the oscillator’s optimal phase, resulting in phase\ concentration (PC) around the optimal phase at on-beat times. In this model, whether a target sound is off-beat, and by how much, can be judged from the phase distance between the phase at target onset and the optimal phase. High PTP would therefore be achieved when oscillator phases are narrowly concentrated around on-beat times. Conversely, low PTP in this model would correspond to a wider phase distribution at on-beat times. We quantified PC using phase-locking value (PLV) and converted it to circular variance (1 – PLV) to obtain a model measure with the same directionality as the behavioral PTP measure, such that lower values indicate greater PTP. Past implementations of oscillatory entrainment treat any input sound as a binary sound onset event, irrespective of its spectral structure. There, the strength of phase-resetting at each event is determined by the entrainment parameter (EP) which reflects the strength of coupling between sensory input and the model oscillator. At EP = 0, cue sounds do not induce phase resetting; as EP increases, cue-driven phase resetting becomes progressively stronger. These models do not capture spectral variation in the input. Hence, they are principally incapable of capturing the observed effects of cue-target match.

To address this shortcoming, we extended the architecture to a bank of spectrally tuned, uncoupled oscillators, each tuned to a certain spectral frequency in the input sound (**Fig. 4A+C**). Each oscillator is phase-reset by a sound only if it contains energy in the oscillator’s preferred frequency band. Phase-locking to a target is then evaluated across all oscillators that are tuned to spectral frequencies contained in the target sound. Intuitively, this model architecture should produce the strongest PLV, that is greatest PTP, when cue and target have identical spectral content, and reduced PLV otherwise (**Fig. 4D**). We quantified the effects of cue and target spectra on model behavior dependent on the mapping of input sound spectrum to phase reset and the effect of cue sound onset dynamics, resulting in three versions of this architecture (**Fig. 4B**). In Model A, the phase-reset by the cue is binary on or off, independently of the sounds’ relative energy content in that band. In Model B, cue input strength is scaled by the respective sound’s energy in the oscillator’s preferred spectral band. In Model C, cue input strength is additionally scaled by the rise time of the cue’s acoustic onset. This last mechanism implements the notion that entrainment may be stronger for sounds with fast rises, as hypothesized in prior literature ^8,14^ and suggested by the cue effects in Experiment 2. Importantly, for all models, we assume that cue and target have identical overall loudness. This avoids potential effects of loudness differences and matches our experimental design, where sound intensities were perceptually equalized for each cue-target combination.

**Fig. 4.**
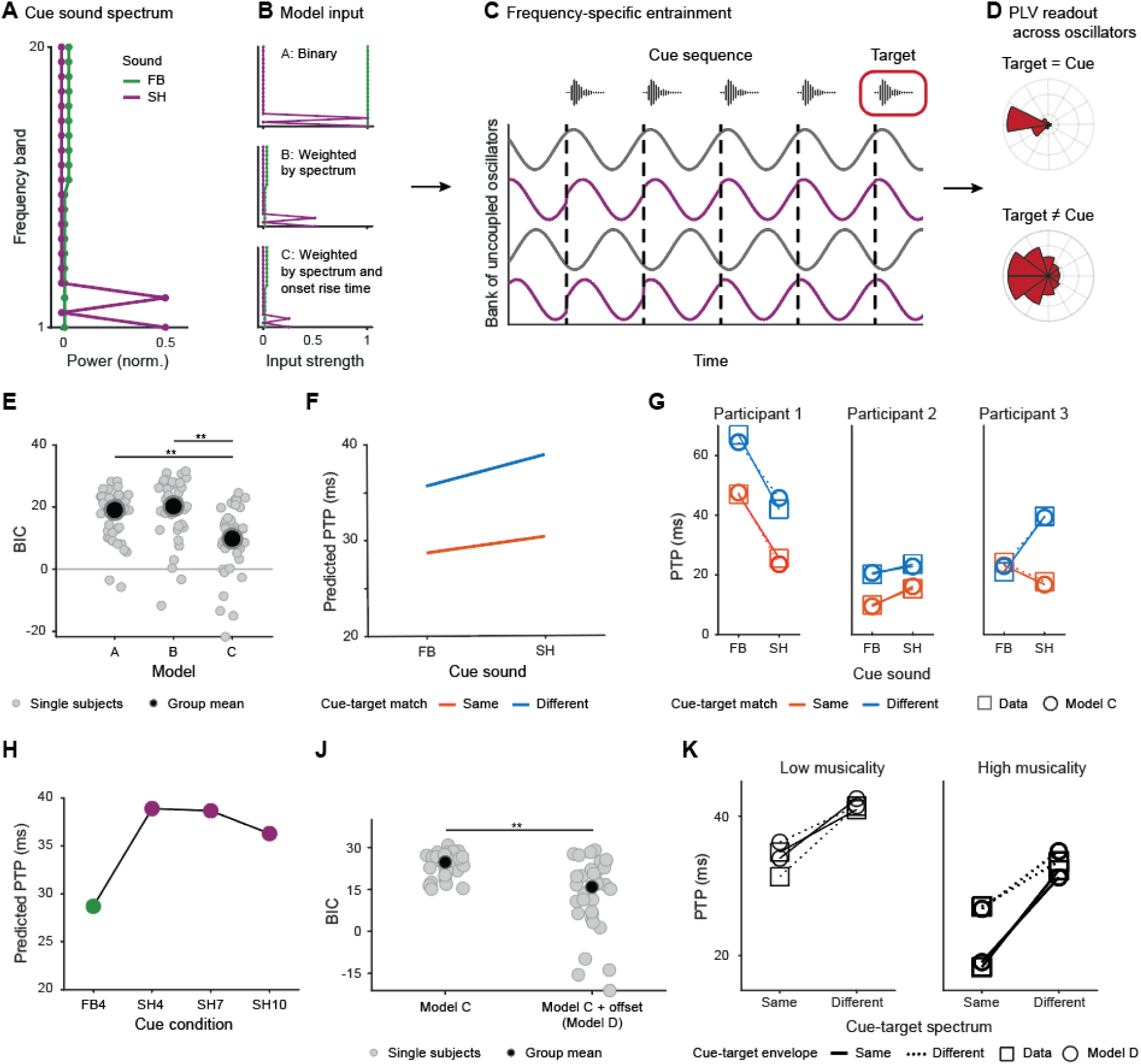
Spectrally-tuned oscillatory entrainment model. **A.** Schematic spectra for fast-rising broadband (FB) and slow-rising harmonic (SH) sound models, per Experiments 1-3. **B.** We modeled three different flavors of the entrainment model. In Model A, input strength for each oscillator was binary, determined as 0 if its preferred spectral frequency was not contained in the cue sound and as 1 otherwise. In Model B, input strength for each oscillator was scaled by the power of its preferred spectral frequency within the cue sound. In Model C, the input strength was additionally scaled down for slow-rising cue onsets. **C.** A bank of uncoupled spectrally-tuned oscillators underwent phase shifts with each cue sound input. **D.** Perceptual timing precision (PTP) was derived from the concentration of phases across oscillators tuned to frequencies contained in the target sound. Phase concentration (PC) was quantified using phase-locking value (PLV), with higher PLV indicating narrower PC. Model predictions were expressed as circular variance (1 − PLV) such that lower values indicate greater PTP. Independently of other model parameters, in all three models PLV was higher when cue and target had the same spectrum than when their spectra differed (here visualized for an example model run). We used a linear transformation to match the scales of model predictions to PTP in the behavioral data. **E.** Model comparison using BIC shows that Model C fits behavior in Experiment 2 better than Models A and B. **F.** Group-level behavior predicted by Model C for Experiment 2 captures the behavioral data, cf. to Fig. 2. Note that this is averaged across predictions for participants in Experiments 2a and 2b. **G.** Data and model predictions for three example participants from Experiments 2a and 2b show that Model C captured a wide range of individual behaviors. **H.** Model C fitted to Experiment 2 also captures the behavior observed in Experiment 1, at the group level. **J.** Model D, an extension of Model C that includes weighted offset entrainment, fits data of Experiment 3 better than Model C. **K.** Model D captures the increase in envelope-match effects for participants with low/high musicality, based on a median split.

To characterize the models’ ability to capture the PTP patterns observed in Experiment 2, we quantified model-predicted PTP as a function of EP and, for Model C, onset-dependent scaling, for all cue-target combinations of a fast-rising broadband sound (FB, modeled with a broad spectrum; **Fig. 4A**) and a slow-rising harmonic sound (SH, modeled with a narrow spectrum with two harmonics for simplicity; **Fig. 4A**). We used a grid-search across a wide range of parameter value combinations to fit each model to individual participants’ data, and calculated BIC for each individual participant based on a linear fit between experimental data and model predictions across the four experimental conditions of Experiment 2 (**Fig. 4E**). While all models were able to partially capture the observed patterns (median BIC Model A: 19.1, Model B: 20.3, Model C: 9.8, **Fig. 4E**), Model C clearly outperformed the other two models (rank-sum test on BIC across participants *p* < .001 for comparison against Model A and Model B). Model C not only accurately reproduced the group-level average behavior in Experiment 2 but was also able to accurately capture a range of individual behavioral patterns (**Fig. 4G**, see **Fig. S2** and **S3** for details on model behavior and fits for individual participants). Notably, the set of Model C parameters that best fit Experiment 2 data (**Fig. 4F**) also replicated the (lack of) length effects observed in Experiment 1 (**Fig. 4H**).

Together, these model comparisons show that capturing the dependence of PTP on spectral continuity between cue and target sounds requires a model that incorporates both spectral tuning of entraining oscillators and cue onset-scaled input strength.

### Offset entrainment captures effects of cue-target envelope continuity on PTP

Our next aim was to extend Model C to capture effects of matching temporal envelope, as observed in Experiment 3 for musically apt participants. For this, we expanded Model C to include information about the temporal dynamics of cue and target sound envelopes (Model D) beyond their onset dynamics. A large body of literature shows that auditory cortices contain not only spectrally tuned onset detectors but also spectrally tuned offset detectors ^36–38^. We therefore introduced a set of offset-entraining oscillators and tested whether varying their contribution to the overall PTP estimate could capture the beneficial effect of matching the temporal envelope between cue and target sounds on PTP. This extended Model D fit the data of Experiment 3 better than an onset-only based Model C (**Fig. 4J**) and accurately captured the increase in envelope-match effects that occurred in participants with higher musicality (**Fig. 4K**). Model D thus provides a potential account of the observed individual variability in benefits from envelope continuity, based on the contribution of sound offset-related temporal information to perceptual timing.

### Greater musical aptitude improves perceptual timing precision

Finally, given the inconsistent effects of musicality across individual experiments, we pooled data across experiments, to test whether beneficial effect of musicality on PTP would depend on cue-target matching (**Table 1**). First, we asked whether benefits from musicality differed between FB–FB and SH– SH sequences. Musicality improved PTP when cues and targets were identical (average musicality slope across FB-FB and SH-SH sequences, *β* = −5.91, 95% CI [−8.34, −3.49], *SE* = 1.23, *t*(151) = −4.82, Holm-adjusted *p* < .001, ηp^2^ = .13; **Fig. 5A**). Although numerically stronger for SH-SH (*β* = −7.09, *SE* = 1.48) than for FB-FB (*β* = −4.74, *SE* = 1.37) sequences, this difference was not significant statistically (*β* = 2.35, 95% CI [−0.54, 5.23], *SE* = 1.46, *t*(164) = 1.61, Holm-adjusted *p* = .110, ηp^2^ = .02; **Fig. 5A**). Second, we tested whether the effect of musicality generalized to sequences in which cues and targets differed in both spectrum and envelope. Musicality improved PTP in these sequences (*β* = −2.90, 95% CI [−5.42, −0.39], *SE* = 1.27, *t*(172) = −2.28, Holm-adjusted *p* = .048, ηp^2^ = .03; **Fig. 5B**), but this effect was smaller than when cues and targets were identical (*β* = −3.01, 95% CI [−5.01, −1.01], *SE* = 1.01, *t*(146) = −2.97, Holm-adjusted *p* = .010, ηp^2^ = .06; **Fig. 5B**). In sum, greater musicality benefits PTP overall, with most benefits under spectro-temporal continuity. This indicates that greater musicality facilitates the use of spectro-temporal continuity between successive sounds.

**Fig. 5.**
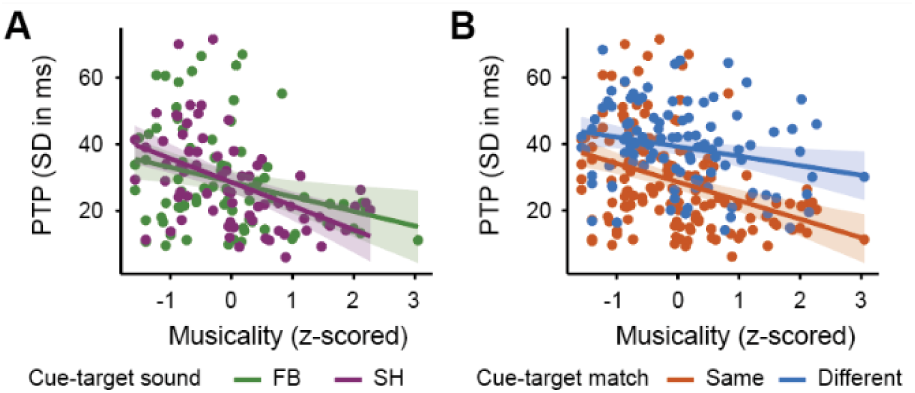
Musicality effects across experiments. **A.** Higher musicality was associated with greater PTP for identical cue-target combinations, both for fast-rising broadband (FB-FB) and slow-rising harmonic (SH-SH) sequences. **B.** The effect of musicality on PTP was weaker, but still significant, when cue and target sounds differed in both spectrum and envelope, than when they were identical.

**Table 1.** Participant counts by experiment and condition in the pooled musicality analysis.

| Experiment/condition | FB-FB | SH-SH | Different |
| --- | --- | --- | --- |
| 1 | 37 (FB4) | - | 35 (SH4) |
| 2a | 21 | 21 | 20 |
| 2b | 19 | 17 | 17 |
| 3 | - | 32 (S=E=) | 31 |
| **total** | **77** | **70** | **103** |

*Note.* Values indicate the number of participants contributing observations to each condition within each experiment. Because participants could contribute to more than one condition, column totals represent participant-condition observations. The pooled model included 250 observations from 111 participants.

## Discussion

We investigated how perceptual timing precision (PTP) in rhythmic contexts is affects by spectro-temporal continuity between acoustics of cues in context sequences and target sounds. PTP was determined by continuity between successive sounds: PTP was best when cue and target were identical. Critically, there were no effects of target acoustics and only minor effects of cue acoustics. A follow up experiment showed that PTP improved most strongly when cue and target had the same spectrum, in line with the spectral specificity of oscillatory entrainment for temporal prediction ^31,32^. Surprisingly, we found that listeners with higher musical aptitude had even better PTP when cue and target also had the same temporal envelope. To account for these findings, we developed an extension of existing oscillatory entrainment models ^26,34,35^, using a bank of spectrally tuned oscillators with separate entrainment to sound onsets and offsets. The model captured key aspects of the behavioral results, hence providing a mechanistic account for the effect of acoustic continuity between successive sounds on the fidelity of temporal predictions and PTP. Overall, our findings show that PTP is not a psychoacoustic property of individual sounds, as suggested in the beat bin account ^8,14^. Rather, it is determined by an interplay of contextual and target sound acoustics, based on neural entrainment of spectrally tuned neural populations.

Most work on temporal prediction mechanisms has focused on the temporal structure of predictive contexts, with less attention to the spectro-temporal dynamics of such contexts ^40^. Our results show that spectro-temporal information is critical for PTP in explicit temporal judgements, providing converging evidence with prior behavioral and electrophysiological work on temporal prediction ^31,32^. Recently, Cheng and Creel ^41^ observed entrainment effects both within and across speech and nonspeech sounds, but found tone-to-tone entrainment to be most reliable. They suggested that greater auditory similarity may facilitate entrainment. Our findings refine this account and identify spectral continuity as the key determinant of PTP. Spectral selectivity is a central property of subcortical and primary auditory regions ^42–47^. Higher-order associative cortical areas, however, exhibit more complex spectro-temporal tuning. Moreover, broader and more heterogeneous frequency tuning is also observed throughout the non-lemniscal subcortical auditory pathway and in posterior superior temporal gyrus ^48–50^. The determining role of spectral selectivity for PTP in rhythmic contexts is therefore consistent with stimulus-driven neural entrainment in frequency-selective auditory populations at primary cortical or subcortical levels.

PTP was also higher in musically apt participants across cue-target combinations. Importantly, the effect of musicality was stronger when cue and target shared spectral content. That is, musicality benefited most in conditions that afforded the highest PTP, suggesting that musicality modulated the continuity effect, rather than more general processing skills. This interpretation is consistent with evidence that musical expertise strengthens cortical tracking and entrainment to auditory rhythms and produces experience-dependent enhancements in auditory feature processing ^51–59^. This effect may be due to improved spectral tuning and hence stronger reliance on spectral overlap with higher musical aptitude. Similarly, PTP was improved, in musically apt participants, when the temporal envelopes of cue and target sounds were identical. Inspired by descriptions of spectrally tuned offset responses in auditory cortices ^36–38^, we hypothesized that this reflects improved neural entrainment of offset populations in auditory cortices and their contribution in perceptual timing judgements.

We integrate our findings in computational models of temporal prediction ^26,34,35^, using a novel spectrally resolved oscillatory entrainment model that captured the effects of spectral and envelope continuity on PTP. Existing computational entrainment models suggest that temporal predictions emerge from neural entrainment to the rhythmic structure of the input ^26^: the phase of endogenous oscillations progressively aligns to rhythmic sensory input such that phases of high neuronal excitability will coincide with predicted time points ^60–62^, leading to improved perceptual sensitivity ^25,27,63^. Entrainment models have been influential in explaining beat and meter perception in music ^26^, including tempo-dependent effects ^64^ and hierarchical metrical organization ^65^. However, current oscillatory modeling approaches are underspecified with respect to how the spectro-temporal structure of individual sounds in a sequence shapes oscillatory entrainment ^40^.

Our modeling approach relies on the same formulation of oscillatory entrainment as used in the past for beat entrainment, modeling perceptual timing precision as the variance in oscillator phase at target time. The key novel aspects of our design were, first, that the input strength for individual oscillators was scaled by both — relative spectral energy in the cue sound and cue onset rise time —, and, second, that model read-out at target time was determined by target spectrum. Together, these parameters captured the entire pattern of our experimental results. Notably, the model parameters that were optimal for Experiment 3 also captured the (absence of) length effects in Experiment 1. In sum, spectrally tuned oscillatory entrainment not only captured the high PTP for sounds with slow-rising onsets, but also reinterprets past reports of PTP dependence on target acoustics as confounding effects of cue-target continuity.

To capture effects of temporal envelope matching, we further stipulated the existence of an offset-entraining population with the same spectral tuning and modeled the effects of musical training via relative weighting of onset and offset phase distributions in the overall phase locking value. The resulting model architecture captures the effects of musicality observed in Experiment 3 and generates several predictions regarding the effects of spectral match and overlap, and different parameters of the temporal envelope for PTP. For instance, future work is necessary to determine whether the effect of musicality reflects enhanced offset-entrainment per se, or whether it facilitates the integration of information across onset and offset entraining populations. Both alternatives can be explicitly modeled and tested against future experimental data to refine the proposed computational architecture. This provides a unified framework integrating known PTP patterns for speech, music and click-like sounds.

Our results can also inform our understanding of temporal predictions in speech. Listeners reliably synchronize to complex continuous signals such as speech and are capable of precise temporal predictions in speech ^19,22,66,67^. In light of our results, we hypothesize that spectral variation within an individual utterance may be sufficiently low to afford high perceptual timing, as it is constrained by articulatory constraints on pitch and formant dynamics ^68–70^. That is, together, these factors may result in sufficiently high spectral continuity to allow for high perceptual temporal precision in speech contexts. It is important to note, that speech is neither isochronous nor rhythmic ^71^ and it is unclear whether oscillatory entrainment contributes to temporal prediction in speech ^35^. However, a distinct body of work on duration judgements for isolated intervals found reduced temporal discrimination when the two sounds marking the interval differed in spectrum ^72,73^. This suggests that our results would also extend to non-rhythmic contexts, even though distinct neural computations might underlie perceptual timing in rhythmic and non-rhythmic contexts.

Our results are also independent of possible interactions between temporal predictions and predictions regarding the identity of a sound. Using distinct experimental designs and non-rhythmic prediction, two recent studies found that expectations about sound rise times ^74^ and their identity ^75^ modulate behavioral and neural measures of temporal prediction. Similarly, Li and Di Luca ^76^ showed that temporal deviations were discriminated more precisely in tone sequences arranged as musical scales than in sequences of randomly ordered tones, irrespective of whether the identity of the final tone was known in advance. They proposed that predictable tonal structure facilitates expectations about upcoming sound frequencies and thereby improves the coding of temporal information. These findings suggest ‘what’ and ‘when’ predictions interact in the auditory system. However, in our study, the (temporal and spectral) identity of the target sound was fully known (due to the blocked experimental design), yet performance was significantly worse when cue and target were not identical. This points to a putative difference between rhythmic-based and symbolic temporal predictions ^34,77^, with sensory entrainment unaffected by expectations regarding the sound identity. The latter may play out at decision or task related performance stages.

## Conclusions

Our results reveal the hitherto disregarded role of spectral and temporal sound contexts for the precision of temporal judgements in auditory perception, refuting the notion that perceptual timing precision is determined by acoustically set sound beat bins. Both were well captured by a spectrally selective oscillatory entrainment model. Cue-target integration provides a powerful and flexible framework for precise temporal analysis of complex sound streams. Overall, we establish spectro-temporal continuity as a key determinant of perceptual timing precision in rhythmic streams and provide a testable mechanistic framework for understanding perceptual timing precision in complex sound streams, such as music and speech.

## Methods

Experiment-specific methods are described in the individual experiment sections.

### Participants

Participants were recruited via a university-wide mailing list and compensated 12€ per hour. Eligibility criteria required participants to be native German speakers, 18-35 years old, with self-reported normal hearing and no history of dyslexia, speech disfluency, or attention deficit disorders. Individuals were not permitted to participate in more than one experiment. All experiments were approved by the ethics committee of the University Hospital Tübingen (209/2022BO2).

### Experimental task

We assessed perceptual timing precision for the target sounds using the Iterative Adjustment Paradigm (IAP; **Fig. 1B**), adapted from Marcus ^16^. In each trial, participants first listened to the stimulus sequence and then adjusted the timing of the target by moving a slider along a horizontal bar on the screen, to indicate when the target should occur to fall on beat with the cue sequence. They then heard the adjusted sequence and were offered to make further adjustments using the same interface. This iterative procedure of listen-adjust-listen was repeated until subjective isochrony was reached, or for a maximum of 12 iterations per trial. Slider movements were continuous, allowing 1 ms temporal resolution in adjustments. The slider was placed in the center of the scale at the beginning of each trial. The IAP was implemented in a custom script in MATLAB R2022b (MathWorks) using Psychtoolbox 3.0.18 ^78,79^.

Participants completed all four IAP conditions in a blocked design, with block order counterbalanced across participants. Each block began with three practice trials to familiarize participants with the stimuli and adjustment procedure. In each condition, participants completed 16 experimental trials (8 with initial positive and 8 with initial negative target offsets). Participants could proceed at their own pace and were encouraged to take breaks between blocks.

### Stimuli

Each trial consisted of a cueing sequence followed by a single target sound (**Fig. 1A**). Cueing sequences contained identical sounds presented isochronously at a rate of 1.5 Hz. The target sound occurred at a pseudo-random temporal offset of 70 −100 ms either before or after the expected on-beat time of the cueing sequence.

To ensure symmetric sampling of early and late targets, we randomly generated positive (late) offset trials for each participant and then mirrored them with negative signs to create corresponding early target trials.

Experiment-specific manipulations of cue and target sound features (e.g., number of cues, complexity, spectrum, or envelope) are described in the individual experiment sections.

### Musical background questionnaire

We assessed musical aptitude with a custom eight-item questionnaire. Items covered the frequency of musical activities, years of formal musical training, current and past musical engagement, and self-rated rhythmic abilities in perception and production (see Table S1 in the Supplementary Materials for the questionnaire items). Responses were pooled across experiments, z-standardized separately for each item, averaged within participants, and the resulting composite score was z-standardized again across participants. Higher scores reflect greater overall musical aptitude, including longer training, and more frequent engagement in musical activities.

### Rhythm discrimination task

In Experiments 2b and 3, we assessed participants’ rhythmic processing abilities using the nonverbal forced choice rhythm discrimination task described by MacIntyre & Scott ^80^. We implemented the publicly made available Gorilla^81^ version of the task and translated the instructions into German. Each trial consisted of a rhythmic pattern of conga beats played twice, followed, after a short pause, by a second rhythmic pattern presented once. Participants indicated whether the second pattern was the same as or different from the first. Each pattern lasted 3.2 seconds and consisted of nine conga drum beats. The task consisted of 40 trials and took approximately 15 minutes to complete.

Rhythm discrimination performance was quantified as the proportion of correct responses for each participant across the 40 trials (accuracy). This score was subsequently z-scored across participants from both Experiments 2b and 3 before being entered in our explanatory analyses.

### Staircase procedure for individual loudness adaptation

In Experiment 3, we implemented an adaptive staircase procedure to control for potential confounding effects of loudness in the IAP. Because perceived loudness depends on spectro-temporal stimulus properties, the four sounds were expected to differ in perceived loudness ^82–84^.

The IAP cue sound (SH) served as the reference, and the other three target sounds were individually adapted relative to it. No adjustment was required for the condition in which cue and target were identical. Participants therefore completed one staircase for each of the three remaining target sounds, with staircase order counterbalanced across participants.

In each trial, participants heard the cue and target sound in random order and indicated which of the two was louder. Based on the response, the level of the target sound was either increased or decreased to converge toward equal perceived loudness.

The staircase followed a 1-up, 1-down procedure with fixed step sizes of 6, 6, 4, 2, 1, 0.5, 0.25, and 0.25 dB. Each staircase terminated after a maximum of eight reversals, and the loudness parameter was calculated as the mean of reversals 5–8. This parameter was then converted into an amplitude scaling factor, which was applied to the target sounds used in the IAP. The procedure was implemented in a custom script (adapted from https://code.google.com/archive/p/matlabstaircase/) in MATLAB R2022b (MathWorks) using Psychtoolbox 3.0.19 ^78,79^.

### Procedure

Testing took place in individual sessions in a quiet room. After providing informed consent, participants completed a demographics questionnaire, followed by the main IAP task, and subsequently a debriefing questionnaire (not analyzed here), a musical background questionnaire and in Experiments 2b and 3 the rhythm discrimination task. The entire session lasted approximately 60-90 minutes. The experiment was administered on a laptop, participants wore over-ear headphones (Beyerdynamics DT990 Pro) connected to an external audio interface (Focusrite Scarlett 4i4 3rd generation), and responded using a USB mouse.

### Data cleaning and preprocessing

Preprocessing of the behavioral data from the iterative adjustment paradigm (IAP) followed the same overall procedure in all experiments. The goal of the preprocessing was to ensure that final adjustments reflected meaningful perceptual corrections, and perceptual timing precision (PTP) estimates were based on a sufficient number of trials.

1. **Exclusion of invalid adjustments.** Our initial offsets were designed to be clearly audible as off the beat. We thus excluded single trials in which the final adjustment resulted in a target offset equal to or larger than the initially presented offset.
2. **Removal of time-out trials.** In Experiment 2a, trials with response times of more than 6 seconds were marked as timed-out and thus removed.
3. **Removal of audio-artifact trials.** In Experiments 1 and 2b, a reproducible audio artifact occurred in a subset of trials due to a scripting error during stimulus playback. The artifact consisted of a truncated final cue sound: specifically, a small portion (up to ∼70-90 ms depending on the experiment) of the end of the final cue tone was cut off. Although some cut-offs were so small that they were not audible, we removed all affected trials.
4. **Minimum trial requirement.** PTP was operationalized as the standard deviation of target offsets after the final adjustment. Of note, to determine PTP no estimate of the perceptual onset location is necessary. To ensure reliable estimates, subject-wise PTP was calculated only for conditions with at least nine valid trials.
5. **Condition-specific outlier removal.** After computing PTP for each participant × condition, outliers were defined as PTP values exceeding ±1.5 × IQR (computed across participants within the same condition) and were removed.
6. **Attention check exclusion.** In Experiment 2a, participants who responded incorrectly to more than half of the attention check catch trials were excluded.
7. **Participant exclusion.** We excluded entire participants if more than half of the experimental conditions were missing after the above steps.

We report the percentage of excluded trials and number of participants separately for each experiment.

### Data analysis

Across all four experiments, we analyzed perceptual timing precision (PTP; standard deviation of responses, in ms) using Linear Mixed-Effects Models (LMMs) implemented in the lme4 package ^85^ in R ^86^. *p* value estimates were obtained from the lmerTest package ^87^. Bayesian modeling was performed using the R BayesFactor package^88^.

### Model selection

Our primary aim across experiments was to evaluate (a) the effects of the experimental manipulations on PTP and (b) whether individual differences in musical aptitude consistently predicted PTP. Musicality was z-scored across all data sets and included as a covariate in all models. All models included a random intercept for participant to account for between-participant variability.

Although rhythm discrimination was also measured next to musicality in Experiments 2b and 3, we did not include it as a covariate in our primary LMMs to maintain a consistent covariate structure across experiments.

### Exploratory analyses of musicality across experiments

To examine whether individual differences in musical aptitude were associated with PTP, we pooled data across all experiments (see **Table 1** for an overview of the included conditions). Specifically, we asked whether (a) musicality improved PTP across identical cue-target sequences and whether this beneficial effect differed between identical sound sequences with different sound properties (FB-FB vs. SH-SH). We further asked whether (b) the beneficial effect of musicality generalized to sequences in which cue and target differed in both spectrum and envelope, and whether its magnitude differed between identical and different cue-target sequences.

The categorical predictor condition (different, FB-FB, SH-SH) was entered in the LMM using treatment coding with the level “different” as the reference level. We included condition, musicality, and their interaction, as well as experiment as a fixed effect and a random intercept for participant nested within experiment.

The final model was:

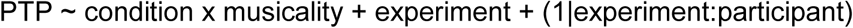

We estimated musicality slopes using estimated marginal trends and conducted four planned tests. To address question (a), we tested the average musicality slope across FB–FB and SH–SH sequences against zero, and compared the musicality slopes between the two identical sequence conditions. To address question (b), we tested the musicality slope for different cue–target sequences against zero, and compared it with the average slope across FB–FB and SH–SH sequences. Holm-adjusted *p* values were used to account for multiple testing across the four planned tests.

### Exploratory analyses of rhythm discrimination

To characterize the role of rhythm discrimination independently of the mixed-effects models, we conducted exploratory participant-level analyses in Experiments 2b and 3. For each participant, mean PTP across conditions was computed, and we examined (a) Pearson correlation between z-scored musicality and z-scored rhythm discrimination, and (b) Pearson correlation between mean PTP and z-scored rhythm discrimination. We further tested in the pooled data of Experiments 2b and 3 whether rhythm discrimination explained variance in PTP beyond musicality using a linear model with predictors for musicality, rhythm discrimination, their interaction, and experiment as a control covariate.

The full model was:

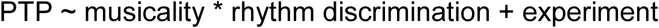

## Experiment 1

### Participants

Thirty-eight participants took part in the experiment. One participant was excluded due to missing data in the musicality questionnaire, resulting in a final sample of 37 participants (29 female, 7 male, 1 diverse; 31 right-handed; *M* = 22.78 years, *SD* = 2.65 years, range 19-30 years).

### Stimuli

Following our hypotheses, we constructed sequences that would allow us to contrast (psycho-)acoustically simple cue tones to more (psycho-)acoustically complex cue tones. We used a fast-rising broadband click (FB, 52 ms duration, broadband spectrum; **Fig. 1C**), resembling a metronome click, as our simple sound. Our complex sound consisted of a slow-rising harmonic tone (SH, symmetrical rise and decay of 150 ms, 300 ms total duration, fundamental frequency 200 Hz, five harmonic components; **Fig. 1C**). In all conditions, the target was the simple sound (FB). In the baseline condition, it was preceded by four simple sound cues (FB4). In the other three conditions, the cue sequence contained four (SH4), seven (SH7) or ten (SH10) complex sound cues.

### Data cleaning

Applying the preprocessing steps described in the General Methods resulted in the exclusion of approximately 16% of trials in total (∼9% due to invalid adjustments and ∼7% due to the audio-playback artifact affecting the SH4 and SH10 conditions). Analyses including these artifact trials yielded the same pattern of statistical results. No participants were excluded based on the data cleaning procedure.

### Data analysis

The categorical predictor condition (four levels) was entered in our LMM using treatment coding with FB4 serving as the reference level.

The full model was:

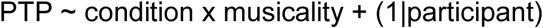

To further assess whether PTP varied systematically with the number of cues within complex cue sequences only, we conducted a post-hoc LMM restricted to the SH-FB conditions. Cue-sequence length was entered as a categorical fixed effect with three levels: short, medium, and long. This allowed us to test whether PTP differed across complex cue-sequence lengths without assuming a linear effect of additional cues.

The model was:

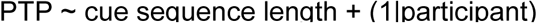

To quantify evidence for the absence of a cue-sequence length effect, we additionally conducted a Bayesian model comparison between the model including cue-sequence length and a null model containing only the participant random intercept. Bayes factors were computed using the JZS prior implemented in the R BayesFactor package ^88^, with the default medium prior scale (*r* = 0.5) for fixed effects.

## Experiment 2a

This experiment’s hypotheses and design were preregistered on OSF. (https://doi.org/10.17605/OSF.IO/M9WQ4). In hindsight, we determined that analyzing the standard deviation of target offsets after the final adjustment would capture PTP best. Consequently, it was not possible to include random slopes in our LMMs. Further, we decided to include musicality as a covariate in all models to maintain a consistent model structure across experiments. A detailed comparison of the preregistered and implemented procedures is provided in Supplementary **Table S2**.

### Participants

Participants were recruited via Prolific ^89^ and received monetary compensation for their participation. Thirty-three participants took part in the experiment. All were English-speaking, aged 18-50 years, and reported normal hearing, and no history of dyslexia, or of speech disfluency. During preprocessing we excluded 11 participants. One further participant had to be excluded due to missing data in the musicality questionnaire, resulting in a final sample of 21 participants (7 female, 13 male, 1 diverse; 18 right-handed; *M* = 28.52 years, *SD* = 6.26 years, range 20-40 years).

### Experimental task

To assess PTP, we implemented the IAP on the online experiment builder platform Gorilla ^81^. The general procedure was unchanged from the other experiments, with minor adaptations due to technical constraints of the platform. In the online implementation, the IAP allowed for a resolution of 5 ms in adjustments, and participants were able to make up to 5 adjustments per trial. We decided to reduce the maximum number of adjustments here as Experiment 1 had shown that participants, on average, only make 2.7 (SD = 1.9) adjustments per trial.

In each IAP condition, participants completed three repetitions of 12 possible initial target onset asynchronies (±70–95 ms), as well as four trials where targets were presented on-beat (overall 40 trials). Due to a technical issue with the online experiment builder, the final trial of each IAP block was frequently not presented to participants. To compensate, single dummy trials were added at the end of each block; these were not included in the analyses. The four trials with zero target-offset were also not included in the analysis of experimental trials, thus each participant could have had a theoretical maximum of 36 experimental trials per condition included in the analysis.

Participants were required to make their adjustment within 6 seconds; otherwise, the trial timed out. To ensure sustained attention and compliance with the instructions, four catch trials were interspersed within each condition block. In these trials, participants heard either a single long tone or three separate tones and had to indicate what they perceived. Three-minute breaks were provided between blocks. The entire session lasted approximately 60–90 minutes.

### Stimuli

The general sequence structure was identical to the other experiments. Here, the target sound occurred at temporal offsets between +-70 to 95 ms in 5 ms increments relative to the expected on-beat time of the cueing sequence.

The experiment followed a 2 (cue sound identity: fast-rising broadband click (FB) vs. slow-rising harmonic tone (SH)) × 2 (cue-target match: same vs. different cue and target sounds) within-subject factorial design, resulting in four cue-target pairings: FB-FB, FB-SH, SH-FB, SH-SH. We used the same two sounds as in Experiment 1: a fast-rising broadband click (FB), and a longer, slow-rising harmonic tone (SH; **Fig. 1C**).

### Procedure

Participants were instructed to complete the experiment in a quiet environment. After providing informed consent, they completed a microphone and headphone screening test, as implemented and made publicly available on Gorilla by Milne et al. ^90^. After instructions and a brief practice session, participants completed the four IAP conditions, which were blocked in counterbalanced order across participants. Following the IAP, participants completed the explicit accelerated version of the Speech-to-Speech Synchronization test ^91^ and afterwards answered demographic, debriefing (not analyzed here) and musicality background questionnaires.

### Data cleaning

Due to data saving issues, the data of four subjects was only partially saved, resulting in a loss of ∼2% of trials. Further ∼4% of trials were affected by random playback errors or timeouts. Together with the preprocessing procedure, this led to the exclusion of approximately 55% of trials. No subjects had to be removed due to failed attention checks. 11 of 33 participants did not meet the minimum-trial requirement (minimum 9 valid trials in at least half of the conditions), and one participant did not complete the musicality questionnaire, they were thus excluded from further analysis. For the remaining 21 participants, an average of 65% of trials was retained after all data cleaning steps, with participant-level retention rates ranging from 35-93%.

### Data analysis

The categorical predictors cue sound identity (FB = 1, SH = −1) and cue-target match (same = 1, different = −1) were entered in the LMM using sum coding.

The final model was:

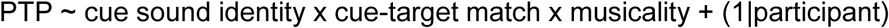

## Experiment 2b

### Participants

Twenty-three participants took part in the experiment. One participant failed to perform the IAP, and another participant did not meet the minimum-trial requirement after preprocessing. These two subjects were excluded from the analysis, resulting in a final sample of 21 participants (15 female, 6 male; 18 right-handed, 3 left-handed; *M* = 22.05 years, *SD* = 2.46 years, range 19-29 years).

### Stimuli

IAP stimulus sequences were identical to Experiment 2a (**Fig. 1C**).

### Procedure

The experimental procedure was the same as described in the general methods section with the addition of the rhythm discrimination task between the debriefing and the musical background questionnaire.

### Data cleaning

Preprocessing removed approximately 20% of trials in total (∼7.5% due to invalid adjustments and ∼12% due to the audio artifact affecting the SH-FB and SH-SH conditions). Analyses including artifact trials produced the same statistical conclusions.

One participant did not meet the minimum-trial requirement (minimum 9 valid trials in at least half of the conditions) and was thus excluded from the analysis. The final sample consisted of N = 21 participants.

### Data analysis

Experiment 2b used the same cue sound identity × cue-target match factorial manipulation as Experiment 2a. Following the unified model-selection strategy described above, only musicality (not rhythm discrimination) was included as a covariate.

The final model was:

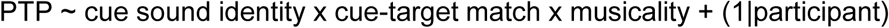

## Experiment 3

### Participants

Thirty-six participants took part in the experiment. Two participants were excluded due to missing data in the rhythm discrimination task, and two additional participants did not meet the minimum required number of trials after preprocessing, resulting in a final sample of 32 participants (26 female, 6 male; 27 right-handed, 4 left-handed, 1 ambidextrous; *M* = 25.72 years, *SD* = 3.72 years, range 20-34 years).

### Experimental task

The implementation of the IAP was run in MATLAB R2022b (MathWorks) using Psychtoolbox 3.0.19 ^78,79^.

### Stimuli

Here, cue sounds were identical across all four conditions and consisted of a slow-rising harmonic tone (SH; **Fig. 3A**) with symmetric rise and decay times (150 ms each, total duration 300 ms; **Fig. 3B**), and four harmonic components starting at a fundamental frequency of 200 Hz (**Fig. 3C**). Note that this SH sound was identical to the SH sound used in Experiments 1 and 2.

The four cue–target pairings were constructed in a 2 x 2 orthogonal manipulation of spectral and temporal envelope match and were as follows.

1. Fully matching condition (S=E=): The target was identical to the above described cue sound (SH; **Fig. 3A-C**).
2. Fully mismatching condition (S≠E≠): The target was a fast-rising broadband click (FB; note that the target is identical to the FB sound used in Experiments 1 and 2; **Fig. 3A-C**).
3. Spectrum-same, envelope-different condition (S=E≠): The target shared the cue’s harmonic spectral content but differed in its temporal envelope. To approximate the fast rise time of the click sound used in the fully mismatching condition, we modeled a target sound with a short rise time (∼2 ms) and a total duration of 107 ms. Due to its harmonic structure, the target’s envelope could not be matched exactly to the metronome click; instead, we implemented a modified version with comparable risetime while avoiding spectral artifacts (FH; **Fig. 3A-C**).
4. Spectrum-different, envelope-same condition (S≠E=): The target preserved the cue’s symmetric 300 ms envelope but employed the same broadband spectrum of the fast-rising click instead of a harmonic one (SB; **Fig. 3A-C**).

### Procedure

The experimental procedure was the same as described in the general methods section with the addition of the staircase procedure before the IAP, and the rhythm discrimination task between the debriefing and the musical background questionnaires.

### Data cleaning

In Experiment 3, preprocessing removed approximately 14% of trials due to invalid final adjustments. No audio artifacts occurred in this experiment. After computing PTP and applying the condition-level outlier and minimum-trial requirements, two participants were excluded for having insufficient numbers of valid trials in more than two conditions. The final sample consisted of N = 32 participants.

### Data analysis

The categorical predictors spectral match (same = 0, different = 1) and temporal envelope match (same = 0, different = 1) were entered in our LMM using treatment coding.

The final model was:

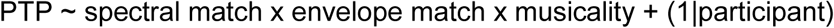

### Oscillator model methods

We implemented a time-resolved coupled oscillator entrainment model, where acoustic sound onsets induce gradual phase corrections in fixed-frequency oscillators. This architecture reflects a typical formulation of oscillatory entrainment models and has been confirmed in previous work ^26,34,35,92,93^. We focused on oscillators whose resonance frequency matched the beat rate of the isochronous cue sequence ^34^. Following Large and Snyder ^26^, we modeled this process using a coupled dynamical system characterized by two state variables, the radius (*r*) and angle (*θ*), as follows:

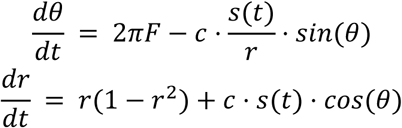

In the absence of external input (s(t) = 0), the system produces periodic limit cycle behavior at a radius of r = 1 and a frequency F. It follows phase correction towards an angle of *θ* = 0 when presented with an input (s(t) > 0). The magnitude of phase correction depends on the strength of the input, the current angle, and the coupling parameter c. To simulate the dynamics of this system, we numerically integrated it using a fourth order Runge-Kutta algorithm with a fixed integration step (MATLAB function ode4.m) and reconstructed the predicted response for each individual oscillator.

Our key goal was to develop a framework to model the impact of spectral content on the entrainment process. For this, we extended the model to a population of spectrally tuned, uncoupled oscillators, each sensitive exclusively to a specific spectral input frequency. We implemented the model with 20 spectral frequency bands, covering the range of audible frequencies between 0.1 and 8kHz. Of note, exploratory simulations confirmed that the simulation results remained qualitatively similar with finer or coarser spectral resolution. For simplicity, we assumed no frequency leakage, namely that each oscillator undergoes phase correction only if the cue sound contains energy at its preferred spectral frequency, and not otherwise. Correspondingly, cue and target sounds were represented as a vector *A*(*f_i_*) at i=1:20 spectral frequency bands. A key assumption of the model is that 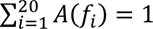, to reflect normalized sound intensity across sounds, warranted by the subjective loudness matching staircase applied in Experiment 3. We modeled the harmonic sounds of our experiments as having equal energy in two frequency bands (for simplicity, results are qualitatively similar with 5 harmonics) and zero elsewhere, and the broadband sounds with low energy in bands at and around the harmonic sounds’ spectral content, and high energy elsewhere.

To translate model dynamics into behavior, we assumed that PTP judgements reflect the degree of phase locking in the system, and modelled it based on phase concentration at target time T across the full population of oscillators. For this, we first calculated the weighted mean phase across all spectrally tuned oscillators for each of N=100 simulations of the entrainment process, where each oscillator had a random initial starting angle between 0 and 2π. We then calculated PLV across all simulations to get an estimate of population-level phase locking. We assumed that PTP for a certain target sound is determined only by the subset of oscillators that are spectrally tuned to frequencies contained in the target sound, with a power above 0. To obtain the overall phase locking value (PLV) across oscillators, each oscillator’s contribution was weighted: 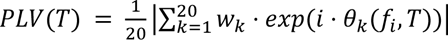, where *θ* (*f*, *T*) is the mean phase of oscillator *f_i_* at target time point T, *w_i_* is 1 for the target sound (*A*(*f_i_*) > 0) and 0 otherwise.

PLV is a measure of phase concentration (values between 0 and 1), with high PLV in this model corresponding to high perceptual timing precision. In contrast, in our experimental work, we estimated PTP as the variability of perceptual timing estimates, such that greater PTP is expressed by lower values. Thus, to obtain a measure of PTP from the model that would have the same directionality as experimentally determined PTP, we use circular variance, defined as *PTP* = 1 − *PLV*(*T*) ^94^.

The magnitude of phase-reset in this model depends on the strength of the environmental input to the model, governed by the entrainment parameter and spectral content of the stimulus. We tested several possible formulations of this model, which differed in conversion of input sounds to model input strength, governing the phase-reset of individual oscillators:

### Model A (binary spectral input)

For each oscillator, the phase-reset by the cue is set to be maximal whenever the cue contains any energy in its tuning frequency, independently of the sounds’ relative energy content in that band (**Fig. 4B** top). In this case, input strength s(*t*, *f_i_*) at cue onset times t, for oscillator *f_i_*, is 1 if cue sound (*A*(*f_i_*) > 0) and 0 otherwise.

### Model B (spectrally-weighted input)

For each oscillator, phase-reset by the cue is scaled by the respective sound’s relative energy in the oscillator’s tuning frequency (**Fig. 4B** middle). In this case, input strength s(*t*, *f_i_*) at cue onset times for oscillator *f_i_* equals to *A*(*f_i_*), the average energy of the cue sound in spectral frequency *f_i_*.

### Model C (spectrally- and onset-weighted input)

We extend Model B such that the magnitude of phase-reset by the cue sound would additionally scale with the risetime of its acoustic onset. This mechanism implements the notion that entrainment may be stronger for sounds with a fast-rising onset, as suggested by the cue effect in Experiment 2b (**Fig. 4B** bottom) and previous reports that onset responses in auditory cortices scale with sound onset risetime ^95,96^. Since our experiment design only contained two types of acoustic onsets, fast- and slow-rising, we assumed a scaling factor of 1 for the fast-rising onset cues and tested a range of multiplicative scaling parameters R, to model phase-correction by the slow-rising sounds: s(*t*, *f_i_*) = *A*(*f_i_*) ∗ *R*.

### Model D (spectrally-, onset- and offset-weighted input)

Models A - C reduce the cue sound to a vector, losing all information regarding its temporal dynamics. They would thus not be able to capture effects of temporal envelope continuity, per Experiment 3. In a second step, we thus expanded Model C (which best captured the data observed in Experiments 1 and 2) to include information about the temporal dynamics of cue and target sound envelopes. We chose to model sensitivity to mismatch in envelope dynamics between cue and target via a second bank of oscillators for which the phase-resetting signal is the offset of the cue sound. This choice was inspired by auditory electrophysiological evidence for frequency-tuned offset responses in auditory cortices ^36–38^. In this model, PLV is calculated across both onset- and offset-entrained oscillators at respective target onset and offset times. If the target’s offset matches the cue’s offset, this contributes additional phases concentrated around the optimal phase. However, if the target’s offset is off-beat relative to cue offset (in case of different envelopes), target offsets contribute phases concentrated around a sub-optimal phase, reducing the overall phase-concentration readout of the system.

To assess model fit, we fitted each model to individual participants’ data using a grid search on a large range of parameter values. Individual model fits were optimized using a linear mapping between predicted and observed PTP values across the four conditions of each experiment. We then used BIC to compare models across participants using a rank sum test.

## Supplementary Material

**Table S1.**

| Item |
| --- |
| Number of years of formal music training (incl. theory and practice) |
| Frequency of professional music making |
| Frequency of rehearsing musical activities |
| Frequency of music making as hobby/amateur |
| Number of musical activities (past and present) |
| Number of years spent on musical activities |
| Average hours per week spent on musical activities |
| Self-assessment of ability to produce a rhythm |
| Self-assessment of ability to judge someone else's rhythm |

*Note.* Musicality Background Questionnaire Items

### Greater rhythm discrimination abilities are associated with higher PTP beyond musicality

**Fig. S1.**
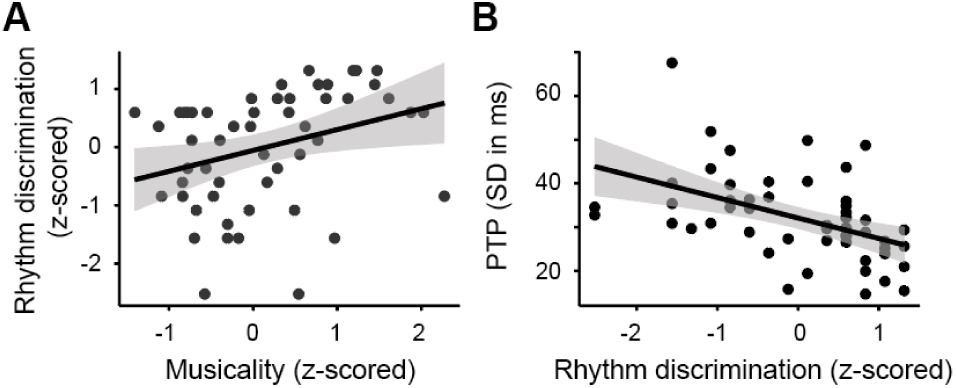
Associations of rhythm discrimination with musicality and perceptual timing precision across Experiments 2b and 3. A+B. Each point represents one participant, lines show linear regression fits, and shaded areas indicate 95% confidence intervals. **A.** Higher rhythm-discrimination scores were associated with higher self-reported musicality. **B.** Perceptual timing precision (PTP), averaged across conditions for each participant, as a function of rhythm-discrimination ability. Higher rhythm-discrimination scores were associated with greater PTP.

The musicality score captures past and present active musical engagement, but may not fully reflect musical aptitude and rhythmic ability. For example, some listeners with little formal musical training may nevertheless have high perceptual sensitivity for detecting rhythmic deviations, whereas musically trained listeners may not necessarily perform equally well on perceptual timing tasks. We therefore also included a rhythm discrimination task in Experiments 2b and 3, as an objective behavioral measure of rhythmic perceptual ability.

Greater rhythm discrimination ability was moderately correlated with higher self-reported musicality (*r* = .32, *p* = .021; **Fig. S1A**). Greater rhythm discrimination was also associated with higher PTP (*r* = −.47, *p* < .001, **Fig. S1B**). To assess whether rhythm discrimination explained variance in PTP beyond musicality, we fit a linear model including musicality, rhythm discrimination, their interaction, and experiment as a control covariate. Both higher musicality and greater rhythm discrimination were independently associated with improved PTP (main effect of musicality, *β* = −2.98, 95% CI [−5.90, −0.05], *SE* = 1.45, *t*(48) = −2.05, *p* = .046, ηp^2^ = .08; main effect of rhythm discrimination, *β* = −3.97, 95% CI [−6.58, −1.37], *SE* = 1.30, *t*(48) = − 3.07, *p* = .004, ηp^2^ = .16). Thus, beyond musical experience, rhythm discrimination captured additional variance in timing precision. This suggests that individual differences in PTP reflect not only accumulated musical training or musical engagement, but also specific perceptual abilities for detecting timing differences. Thus, the ability to evaluate target timing relative to a preceding rhythmic context appears to, at least in part, be associated with listeners’ general sensitivity to rhythmic temporal structure.

**Table S2.**
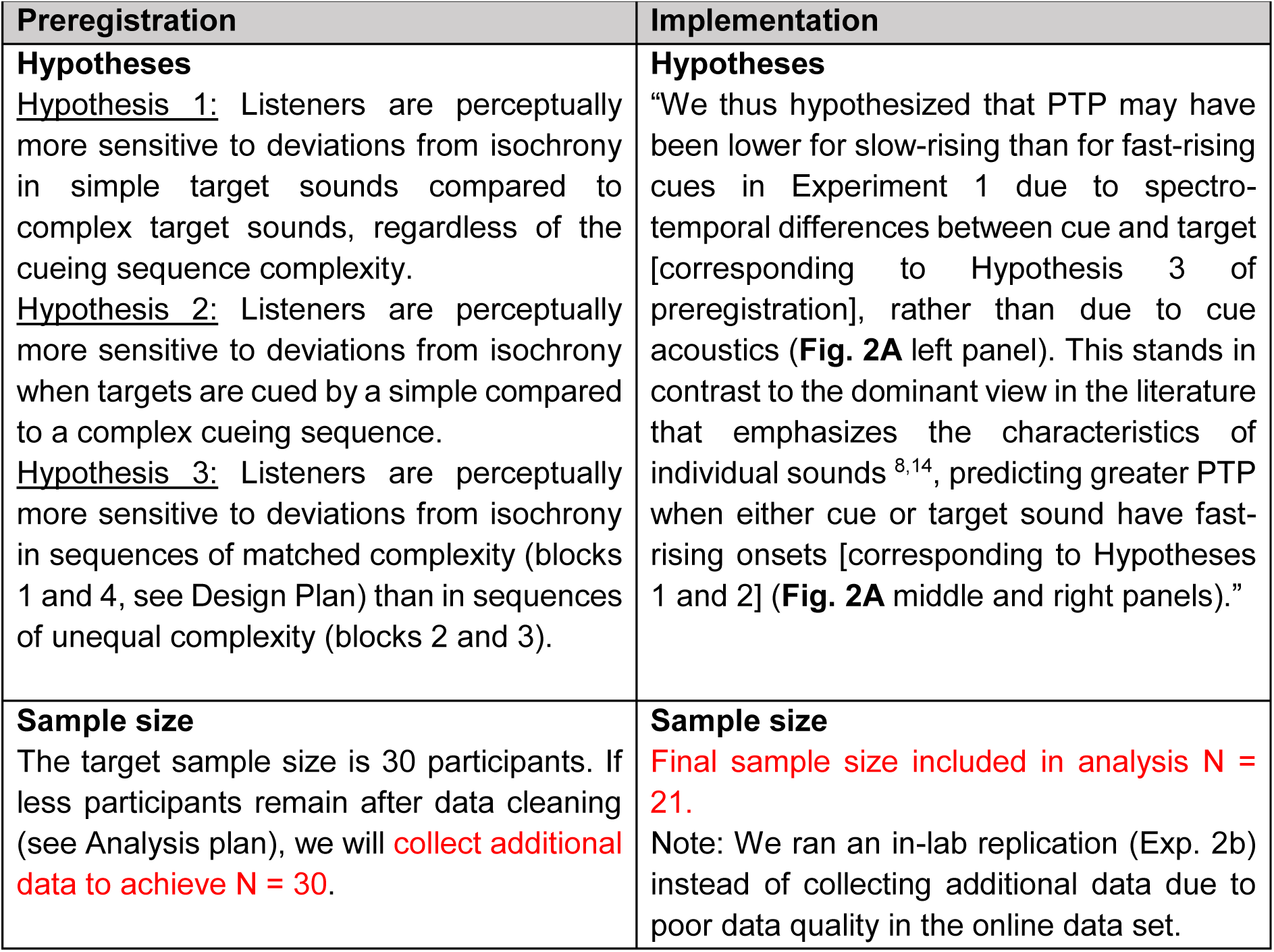
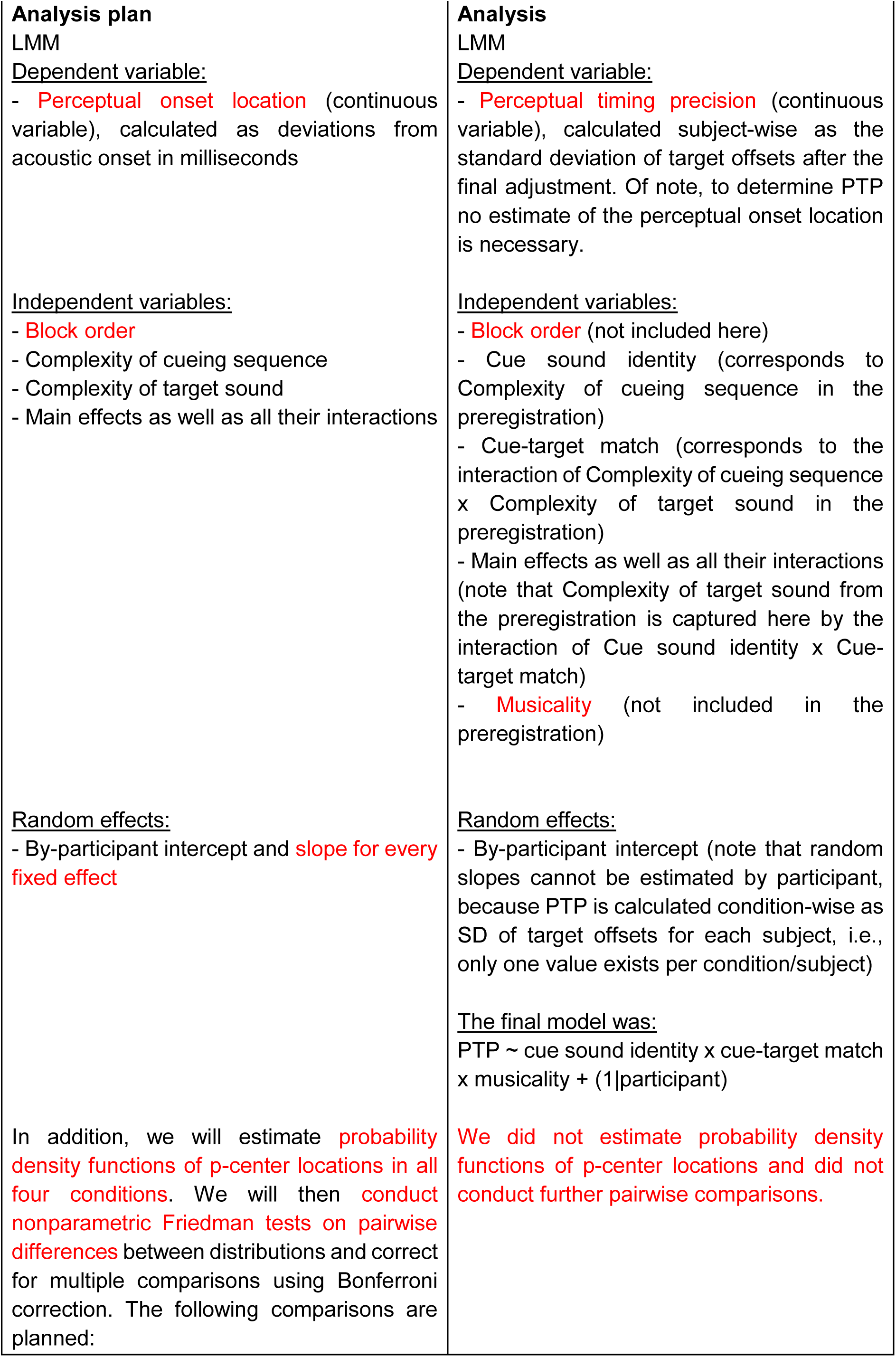
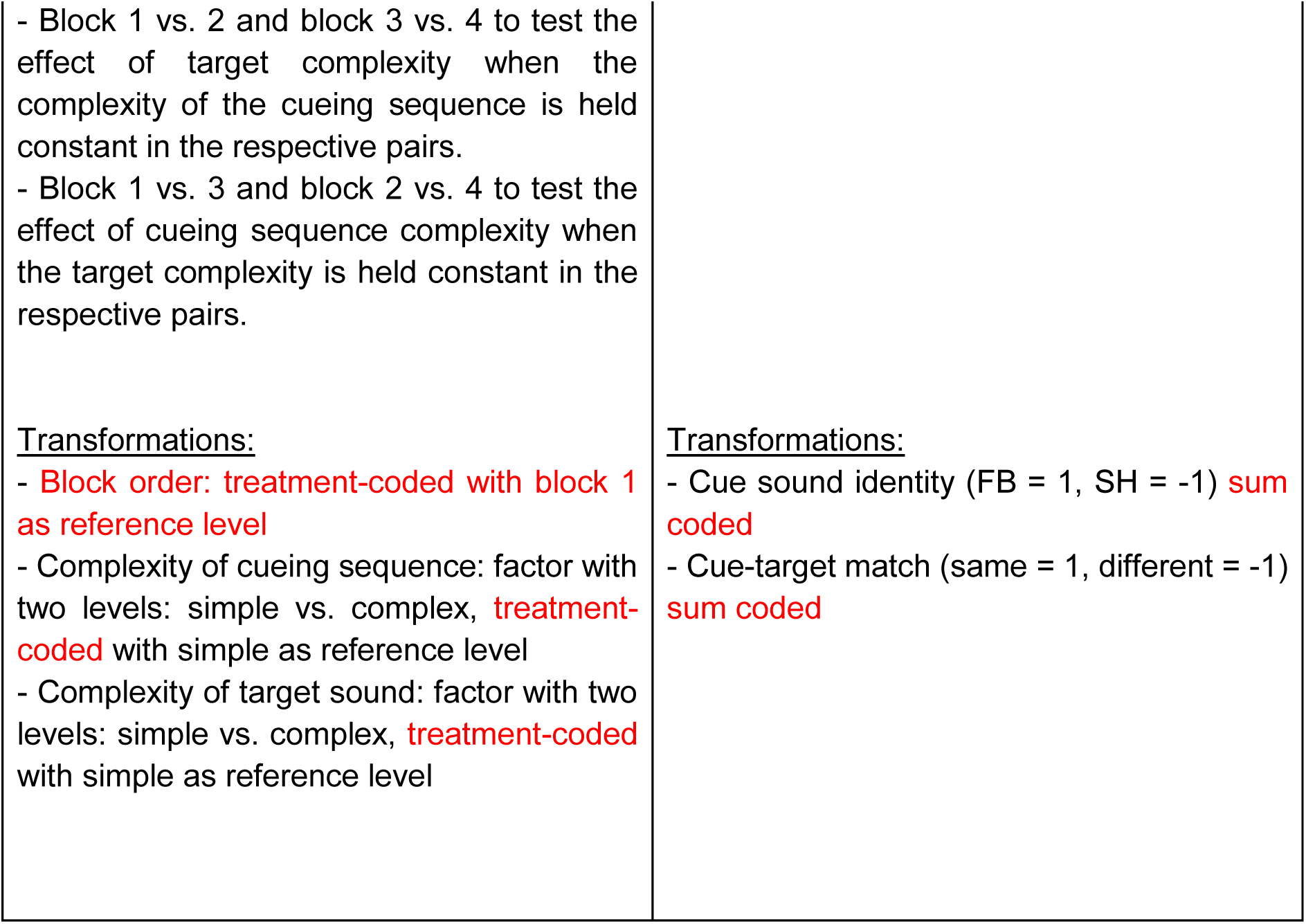
This overview compares the preregistered hypotheses and analyses (https://doi.org/10.17605/OSF.IO/M9WQ4) to the implemented analysis of Experiment 2a. Deviations from the preregistration are highlighted in red.

**Fig. S2.**
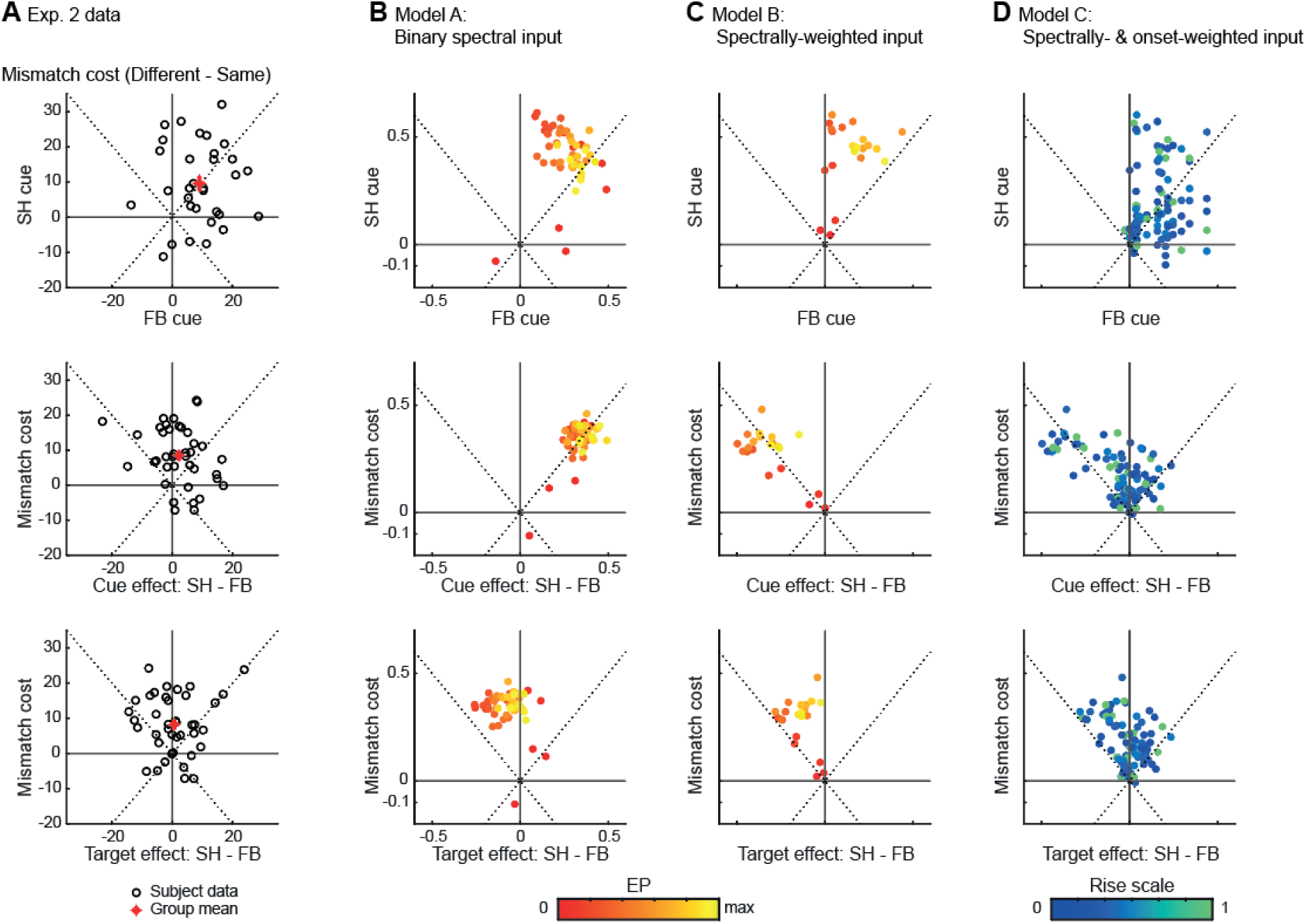
Model behavior. **A.** Distribution of individual behavioral data from Experiments 2a and 2b. Top: Mismatch Cost for a fast-rising broadband cue (FB) versus a slow-rising harmonic (SH) cue. Most participants had higher PTP for matching cue-target pairs. The relative magnitudes of Mismatch Cost for FB and SH sequences were not correlated. Middle: Mismatch Cost versus main effect of cue. Bottom: Mismatch Cost versus main effect of target. Participants were overall better for FB cues than for SH cues, however the effect of cue sound was smaller in magnitude than Mismatch Cost. Target effects varied across participants, with no significant difference between targets at the group level. **B-D.** Same as A for each of the three model versions, as function of entrainment parameter (for Models A-B) and rise-based scaling (for Model C). Only Model C captures the wide range of values and effect patterns observed in the data.

To further understand the models’ behavior, we calculated the three effects that characterized participants’ behavior in Experiment 2:

1. *Mismatch Cost* separately as PTP(cue ≠ target) - PTP(cue = target). Mismatch Cost captures the detrimental effect of mismatch between cue and target sounds. We calculate it once separately for fast-rising broadband (FB) cues and slow-rising harmonic (SH) cues, and once averaged across both types of cues, effectively modeling the main effect and both simple effects of cue-target match.
2. *Cue Effect* as the difference between conditions with FB cues and SH cues (PTP(*FB*- FB) + PTP(*FB*-SH) - PTP(*SH*-SH) + PTP(*SH*-FB)). In Experiment 2a, this effect was non-significant, whereas in Experiment 2b PTP was a bit better for FB cues than for SH cues.
3. *Target Effect* as the difference between conditions with FB targets and SH targets (PTP(FB-*FB*) + PTP(SH-*FB*) -PTP(SH-*SH*) -PTP(FB-*SH*)). This effect was not significant in both Experiments 2a and 2b.

We then visually assessed the distribution of PTP patterns generated by each model and compared it to individual participants’ behavior observed in Experiment 2 (**Fig. S2A**). A model that captures participants’ behavior in Experiment 2, should be able to produce a range of similarly large Mismatch Costs for FB and SH cues (**Fig. S2A top**), a positive cue effect of smaller magnitude than the Mismatch Cost (**Fig. S2A** middle), and a range of target effects around 0 (**Fig. S2A** bottom). Importantly, cue and target effect magnitudes should be independent of Mismatch Cost magnitude, as these effects were not correlated in our behavioral data (*r* = 0.1, *p* > 0.1). As expected, all three models captured the Mismatch Cost between cue and target (**Fig. S2B-D** top row) to some degree, with all data points across parameter values in the upper right quadrant. However, neither Model A nor Model B could capture the full pattern of results. First, only Model C was able to produce a similarly broad range of Mismatch Cost for SH and FB cues as observed in the data, whereas Models A and B underestimated the effect for SH cues (**Fig. S2B-D** top). Second, in Models A and B, the cue effect was strongly correlated with the Mismatch Cost (**Fig. S2A B-D** middle row). Moreover, both models predicted a negative target effect for most values of EP (**Fig. S2B-D** bottom row). Both qualitatively diverge from the experimental data. In contrast, introducing onset-based scaling of input strength in Model C alleviated this coupling and generated a wider range of values for Mismatch Cost, cue effects, and target effects (**Fig. S2D** middle and bottom rows). This analysis is in line with the lowest BIC scores for Model C and further supported by the best match between observed and predicted effects for Model C (**Fig. S3**).

**Figure S3.**
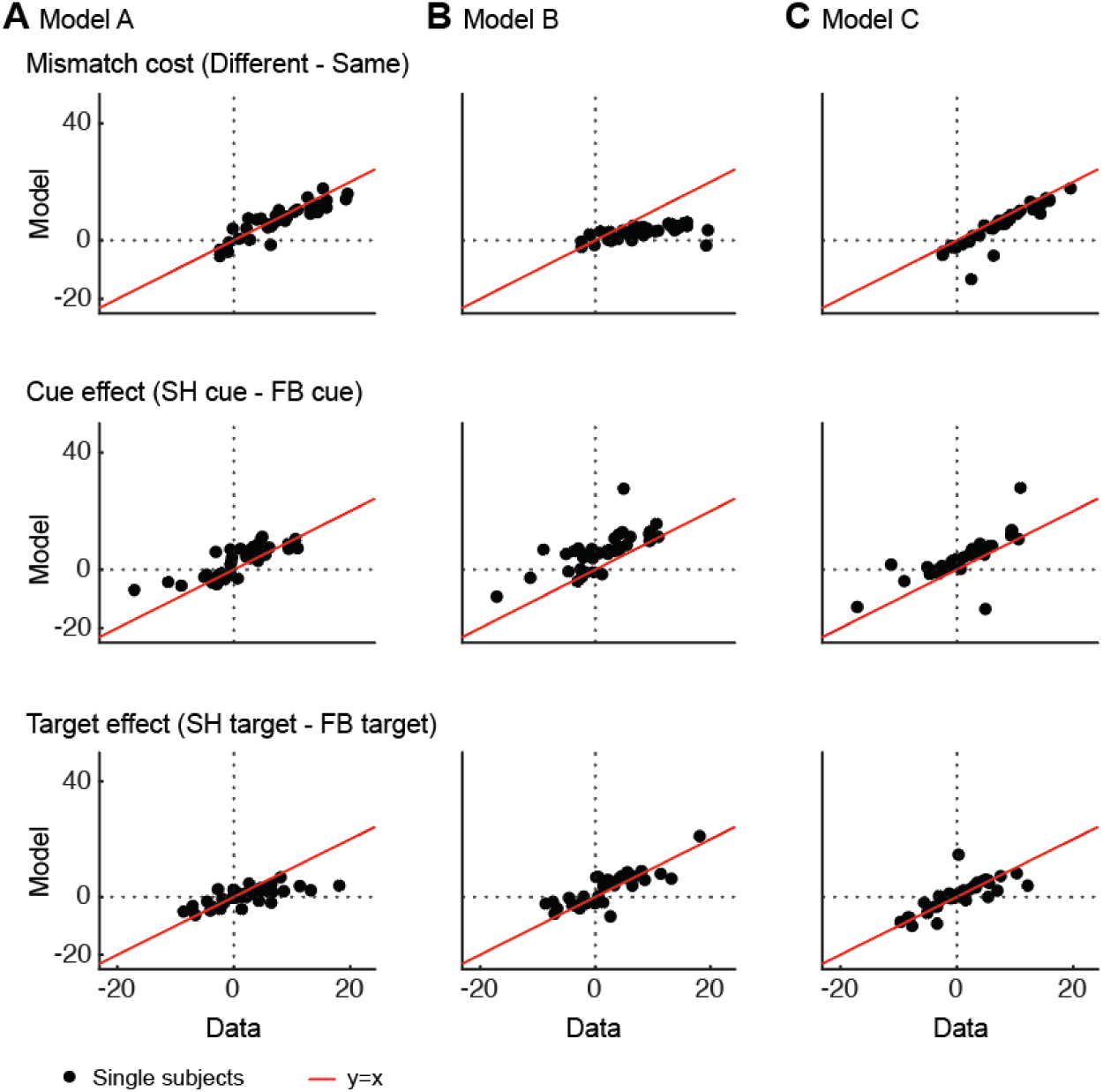
Model fit to Experiment 2 data. **A.** Model A: Binary spectral input. **B.** Model B: Spectrally-weighted input. **C.** Model C: Spectrally- and onset-weighted input. Only Model C captures the wide range of values and effect patterns observed in the data.

